# Identification of the Best Filter-based Feature Selection Techniques for Microarray Datasets

**DOI:** 10.1101/2025.04.25.650719

**Authors:** Atikul Islam, Tapas Bhadra, Kalyani Mali, Jayant Giri, Khursheed Aurangzeb, Ram Kaji Budhathoki, Saurav Mallik

**Author notes:** Corresponding Authors: Ram Kaji Budhathoki; Saurav Mallik.

## Abstract

This article basically explores the impact of univariate & multivariate filter-based feature selection methodologies on enhancing the classification performance for the real-life classification problems. Our study considers two univariate filter-based feature selection techniques, namely, *Chi-square* and *Fisher score*, as well as two multivariate filter-based feature selection techniques, viz., *Symmetrical Uncertainty* and *Minimum Redundancy-Maximum Relevance (mRMR)*. These methods are applied to feature selection from Five diverse collections of datasets, including datasets related to *Mixed-lineage Leukaemia (MLL), Lung Cancer, Ovarian Cancer, Central Nervous System (CNS)*, and *Colon Cancer*. For each feature, fitness values are calculated using the four aforementioned feature selection methods. After that, a stratified 10-fold cross-validation procedure is conducted using Support Vector Machines (SVM) and Multilayer Perceptron’s (MLP) to determine the classification accuracy for each feature. A set of five microarray datasets was used in this evaluation in order to assess the effectiveness of the filter methods. The results of this study represent the first comprehensive analysis and comparison of gene expression datasets filtered using a variety of ranking strategies. Among these approaches, entropy-based methods (e.g., mRMR) emerge as the most effective. The mRMR method demonstrates Outstanding performance outcomes of accuracy, F1-score, and Root Mean Square Error (RMSE). When comparing classifier performance, the F1-score, which combines precision and recall, is particularly useful, while the RMSE measures prediction accuracy. Chi-square, Fisher Score, and Symmetrical Uncertainty (SU) follow as the second, third, and fourth best approaches, respectively. Although the SVM classifier demonstrates superior performance, the difference in accuracy between SVM and the MLP classifier is marginal.

**Key Points:**

- Our study considers two univariate filter-based feature selection techniques, namely, *Chi-square* and *Fisher score*, as well as two multivariate filter-based feature selection techniques, viz., *Symmetrical Uncertainty* and *Minimum Redundancy-Maximum Relevance (mRMR)*.
- These methods are applied to feature selection from Five diverse collections of datasets, including datasets related to *Mixed-lineage Leukaemia (MLL), Lung Cancer, Ovarian Cancer, Central Nervous System (CNS)*, and *Colon Cancer*.
- After that, a stratified 10-fold cross-validation procedure is conducted using Support Vector Machines (SVM) and Multilayer Perceptron’s (MLP) to determine the classification accuracy for each feature.
- Among these approaches, entropy-based methods (e.g., mRMR) emerge as the most effective. The mRMR method demonstrates Outstanding performance outcomes of accuracy, F1-score, and Root Mean Square Error (RMSE).

## 1. Introduction

In the past few decades, the emergence of DNA microarray datasets has ushered in a dynamic research landscape at the intersection of bioinformatics and machine learning (*Bolon et al, 2014*) [1]. These datasets offer a powerful means to glean critical insights into gene expression variations across diverse tissues and cell samples, with applications spanning from disease diagnosis to tumor classification (*Lazar et al, 2012*) [2]. It is important to note that, due to resource limitations and high costs, these datasets are often limited in terms of the number of genes, or features, which can be effectively included, possibly impeding their application for classification (*Kohavi and John, 1997*) [3].

The advent of DNA microarrays, often referred to as DNA chips, has revolutionized biological experimentation by enabling the simultaneous measurement of gene levels of expression for hundreds of genes, fundamentally transforming our understanding of cellular processes (*Hanai et al, 2006*) [4]. Yet, the reliability and accuracy of subsequent analyses hinge significantly on the normalization process. Normalization contributes significantly to mitigating various sources of variation that may skew gene expression measurements, ensuring that systematic experimental biases and technical fluctuations are properly accounted for and corrected (*Park et al, 2003*) [5]. This indispensable preprocessing step underpins the integrity of subsequent analyses, making it a focal point in the study of microarray data (*Liu et al, 2019*) [6].

Normalization methods encompass various techniques such as scaling and mapping, aimed at levelling the statistical playing field, where each feature contributes equitably to downstream analyses (*Singh and Singh, 2019*) [7]. Scaling and preprocessing are integral aspects of this normalization process, systematically preparing the data for robust analysis (*Patro and Sahu, 2015*) [8].

In the realm of predictive modelling, feature selection is a critical practice employed to streamline input variables, thus potentially enhancing computational efficiency and model performance (*Hira et al, 2015*) [9]. Univariate feature selection methods have been widely explored to improve multi-gene survival predictors, with the Chi-square and Fisher score serving as prominent examples in our study (*Emura et al, 2018*) [10]. The Chi-square statistic quantifies the deviation of observed distributions from expected distributions, assuming feature-class independence (*Sumaiya et al, 2019*) [11]. This approach has shown promise in enhancing classifier performance across various applications (*Bahassine et al, 2018*) [12]. Meanwhile, the Fisher score focuses on identifying feature subsets that maximize the distinction between classes while ensuring compactness within each class, effectively reducing data dimensionality (*Sun et al, 2021* [13], *Quanquan et al, 2012* [14]).

In addition to univariate methods, we delve into multivariate feature selection techniques, specifically Symmetrical Uncertainty (SU) and Maximum Relevance Minimum Redundancy (mRMR) (*Sun et al, 2015*) [15]. SU evaluates the relevance between features and class labels, emphasizing filter strategies that assess individual feature quality and ranking (*Kannan and Ramaraj, 2010* [16], and *Lin et al, 2019* [17]). Similarly, mRMR prioritizes features based on their relevance to the target variable while minimizing redundancy, striking an optimal balance between mRMR (*Li et al, 2012*) [20].

In our study, we applied these four feature selection methods to microarray data, ranking the features based on their respective scores. Following that, we generate classification models using Support Vector Machines (SVMs) and Multi-Layer Perceptron’s (MLPs). Support Vector Machines, a robust supervised machine learning technique, have demonstrated remarkable versatility across a spectrum of biological analysis tasks. These tasks encompass microarray expression data analysis, far reaching protein homology detection, and precision recognition of translation initiation sites (*Furey et al, 2000*) [21]. SVMs shine in both regression and classification scenarios, with the “hyperparameter” C governing margin constraint violations in classification settings (*Gold and Sollich, 2003*) [22]. Drawing from statistical learning theory, SVMs have garnered significant attention for their robust performance and high predictive accuracy (*Zhang et al, 2012*) [23].

On the other hand, Multi-Layer Perceptron’s (MLPs) relies on cross-validation techniques A cross-validation that eliminates one gene and leaves four genes out to uncover diagnostic genes that can be used to analyze diseases (*Zhang et al, 2008*) [24]. To synthesize, our study embarks on an exploration of various feature selection methods, spanning both univariate and multivariate domains, as applied to microarray datasets in the context of classification tasks. Through comprehensive evaluation, we shed light on the efficacy of these techniques in enhancing classification performance in the domain of gene expression data analysis.

## 2. Related Work

According to *Khalsan et al (2023)* [25], a FGS (fuzzy gene selection) method had been designed to identify Insightful genes while reducing dimensionality. This method assesses three feature selection techniques, viz., Mutual Information, Chi-squared, and F-Classif to prioritize genes according to their importance. It employs fuzzification and defuzzification processes to derive an optimal score for each gene, refining the feature selection process. The effectiveness of the FGS method was tested on six gene expression datasets, where it surpassed traditional selection techniques with higher accuracy, recall, precision, and F1-score. This research underscores the potential of integrating fuzzy logic with statistical techniques to enhance feature selection for cancer classification models.

According to *Khan et al (2024)* [26], an improved selection of discriminative features for class-imbalanced gene expression datasets is achieved using a novel robust weighted score for unbalanced data (ROWSU). Synthetic data points are created for the minority class to address class imbalance, resulting in a more balanced dataset. Subsequently, a greedy search strategy is employed to select an efficient subset of genes. The feature selection process is further refined with the addition of a unique weighted robust score, utilizing support vectors. A comprehensive evaluation of the effectiveness of this method was conducted across six gene expression datasets, and k-nearest neighbors (kNN) and random forest (RF) classification techniques were compared to its results. The purpose of this study was to enhance gene expression classification accuracy through class balancing and robust scoring mechanisms.

*Nematzadeh et al (2025)* [27] introduces a statistically based filter approach that evaluates gene significance by considering effective ranges, aiming to improve classification accuracy. Using six benchmark gene expression datasets, FSAER outperformed traditional filtering methods, enhancing SVMs, Naive Bayes, and K-Nearest Neighbors (k-NN).

Key observations drawn from the literature survey are outlined below:

- The FS method has been used in numerous ways, as it plays an important role in determining the most important aspects before classification of the data set is performed.
- For microarray data classification, most authors used an SVM classifier; hybrid approaches were used even more frequently.
- Researchers have used information related to leukaemia, ovarian and breast cancer among the data sets available in the literature survey.

Several researchers have explored various feature selection techniques combined with different classifiers to enhance classification performance. *Osareh and Shadgar (2010)* [28] employed the Signal-to-Noise Ratio (SNR) method for feature selection and tested their approach with Support Vector Machines (SVM), k-Nearest Neighbors (KNN), and Probabilistic Neural Networks (PNN).

According to *Salem et al (2011) [29], a* novel filter-based gene selection method, MGS-CM, was introduced and combined with SVM, KNN, and LDA classifiers to form MGS-SVM, MGS-KNN, and MGS-LDA systems. These models were validated on three microarray datasets, achieving up to 100% accuracy in leukemia classification and minimal misclassification in lymphoma. The approach effectively reduced dataset size by over 99.6% while maintaining high classification reliability. According to *Wang et al (2007) [30]*. Introduced to extract minimal yet highly informative gene subsets for cancer classification from microarray datasets using supervised learning techniques. This strategy effectively lowers computational load and diagnostic expenses while offering insights into gene-cancer relationships. Remarkable accuracy was attained using just 2–3 genes for simpler datasets and 28 genes for a more complex dataset involving 14 cancer types through a divide-and-conquer method. The results highlight the potential for precise diagnosis with a significantly reduced number of genes.

In another study by *Zhang et al (2007) [31]*, Based Bayes error Filter (BBF) was proposed to enhance microarray data classification by removing redundant genes and selecting the most relevant ones. Experimental results on five datasets showed improved classification accuracy with smaller, more efficient gene sets. In *Hang et al (2008) [32]*, data mining has emerged as a valuable tool in cancer research, enabling the discovery of patterns and relationships within large medical datasets to improve disease diagnosis and prognosis. Techniques such as KNN, Naïve Bayes, and SVM are widely applied to analyze cancer data and support clinical decision-making. In *Bharathi and Natarajan (2010) [33]*, a Kernel Fuzzy Inference System (K-FIS) with t-test feature selection was utilized to classify leukemia microarray data by transforming it into a higher-dimensional space using kernel functions. Performance comparison with Support Vector Machine (SVM) revealed that K-FIS delivered similar results across various evaluation metrics, demonstrating its effectiveness and consistency. *Tang et al (2010) [34]* explores the application of gene expression data for cancer classification using Discriminant Kernel-PLS, a machine learning approach that integrates kernel techniques with partial least squares (PLS) to effectively distinguish between various cancer types. In another study by *Furey et al (2000) [35]*, a Support Vector Machine (SVM)-based approach was developed for analyzing DNA microarray data to classify tissue samples and detect potential mislabelling. Applied to ovarian cancer datasets, the method achieved accurate classification and identified key genes, showing comparable performance with other machine learning techniques on additional datasets. In *Gayon et al (2002) [36]*, Support Vector Machine (SVM) learning, known for maximizing the separating margin, has become a key tool in cancer genomics for classification and biomarker discovery. With the rise of high-throughput data, SVMs continue to play a vital role in identifying drug targets and understanding cancer-driving genes. As per *Ye et al (2004) [37]*, Uncorrelated Linear Discriminant Analysis (ULDA) has been proposed to improve classification of undersampled gene expression data by reducing dimensionality through Generalized Singular Value Decomposition, showing superior performance over existing methods. Similarly, *Liu et al (2005) [38]* proposed a Bayesian Network (BN) model was developed to enhance the reliability of prostate cancer biomarkers by integrating data from various biological sources. Both approaches contribute to more accurate and biologically meaningful cancer diagnosis by addressing high dimensionality and data integration challenges. *Yu and Liu (2004) [39]* proposed a method focused on reducing redundancy in feature selection to enhance the effectiveness of microarray data analysis, leading to improved classification and lower dimensionality. In a related study, *Peng et al (2007) [40]* developed a hybrid technique for identifying biomarkers from microarray gene expression data, achieving better performance in classifying cancer types. Overall, this collection of studies highlights the diversity of feature selection techniques and classification models applied across different research works, underlining the ongoing efforts to improve the accuracy and efficiency of classification systems.

In this current study, four feature selection (FS) methods, viz., Chi-square, Fisher Score, Symmetrical Uncertainty (SU), and Minimum Redundancy Maximum Relevance (mRMR) are applied in combination with two machine learning algorithms, such as Support Vector Machine (SVM) and Multi-Layer Perceptron (MLP). SVM and MLP are utilized to classify samples into cancerous and non-cancerous categories for the MLL and colon cancer datasets.

## 3. PROPOSED METHOD

In this paper, an analysis of three-category objects was conducted using a strategy to compare multivariate and univariate feature selection data processing. The regular normalization filter is also applied, as are feature selection methods entirely based on (univariate) chi-square and Fisher scores, as well as (multivariate) SU and mRMR feature selection searchlights. We compared the prediction accuracy of SVM, MLP, and their combination, which have been proven effective in handling high-dimensional data. In this work, firstly take microarray datasets, this data preprocessing in filter based technique normalization. This data is normalized because any information takes meaning full data. Next step we will use the feature selection methods, they are two sorts of feature selection methods are taking univariate feature selection, such as Chi-square and fisher score feature selection method and multivariate feature selection methods such as symmetrical uncertainty (SU) and mRMR feature selection method. After that, we will sort the feature in decreasing order according to the feature score. Next, the proposed method chooses the most informative function from the sorted array. After that used the classifier support vector machine (SVM) and Multi-layer proception (MLP) then the check for the minimal number of features and maximum accuracy. At last, we will calculate the fantastic feature selection methods.

### 3.1 Min-Max Normalization

Min-max normalization is a widely used technique where the minimum value of a feature is scaled to 0, the maximum value to 1, and all other values are mapped proportionally within the range of 0 to 1 (*Liu et al, 2019)* [41]. For instance, if a feature’s minimum and maximum values are 20 and 40 respectively, a value of 30 would be normalized to approximately 0.5, as it lies midway between them. The normalization formula is given below (*Liu et al, 2019) [41], Singh et al*, 2019 [42]).

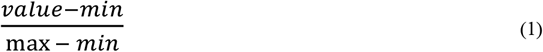

### 3.2 Used Datasets

A microarray database is a repository containing microarray gene expression data. The key uses of a microarray database are to save the measurement data, manipulate a searchable index, and make the records handy for different purposes for analysis and interpretation and better classification accuracy (*Islam et al, 2024)* [43]. Here we are taking two different microarray datasets (https://csse.szu.edu.cn/staff/zhuzx/datasets.html) (*Li et al, 2024)* [44]. The Mixed-Lineage Leukaemia (MLL) dataset consists of 12,582 features (genes) and 72 samples, categorized into three classes. The Central Nervous System (CNS) dataset contains 7,129 features and 60 samples, divided into two classes. The Lung Cancer dataset comprises 12,533 features across 181 samples, categorized into two classes as well. The Ovarian Cancer dataset includes 15,154 features and 253 samples, classified into two categories. Finally, the Colon Cancer dataset contains 2,000 features and 62 samples, divided into two classes: “cancer” and “normal.” The classifiers are executed sequentially on varying feature set sizes [2, 100] across these microarray datasets to evaluate their performance (*Zexuan et al, 2007 [45]* and *Golub et al, 1999)* [46]). The dataset description is the given in **Table 1**.

**Table 1.**
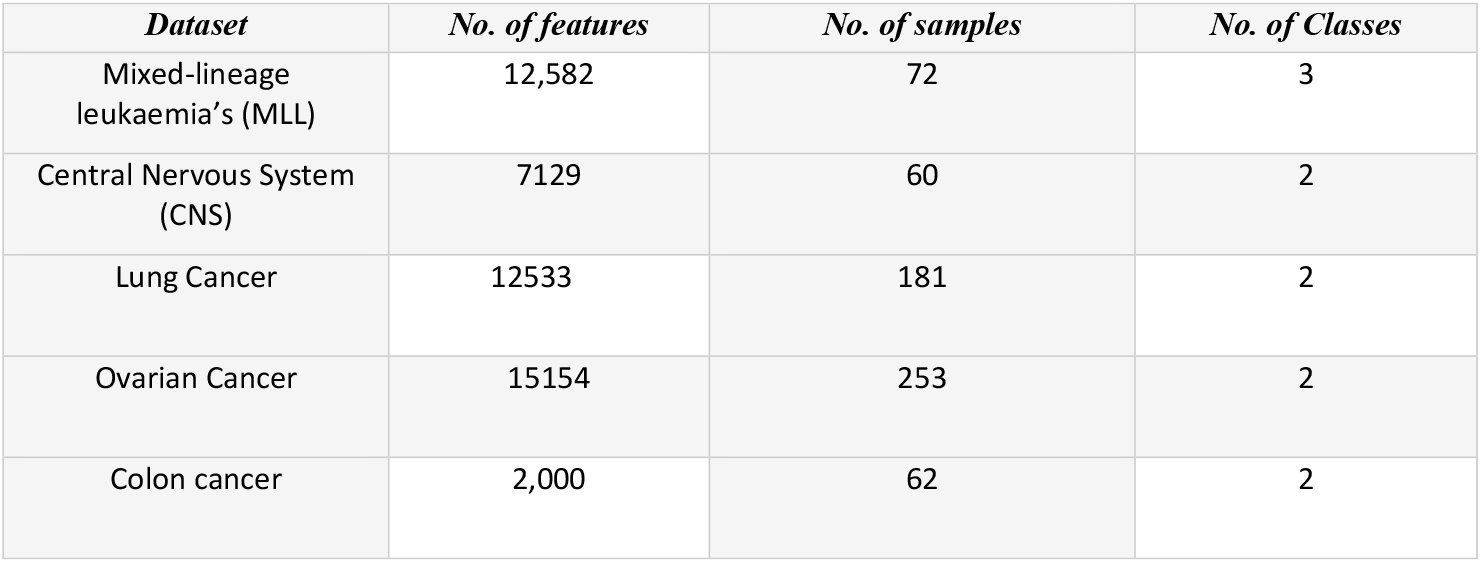
Details of the datasets.

| *Dataset* | *No. of features* | *No. of samples* | *No. of Classes* |
| --- | --- | --- | --- |
| Mixed-lineage leukaemia's (MLL) | 12,582 | 72 | 3 |
| Central Nervous System (CNS) | 7129 | 60 | 2 |
| Lung Cancer | 12533 | 181 | 2 |
| Ovarian Cancer | 15154 | 253 | 2 |
| Colon cancer | 2,000 | 62 | 2 |

### 3.3 Used Feature Selection

#### 3.3.1 Chi-Square feature selection

The chi-squared statistic is employed to compute feature weights, which are then used to establish the ranking of the features. The chi-squared test is performed on both nominal and numerical values. In the first place, it is used to discretize the values of the numerical features, according to a variety of optional discretization techniques using the function Process Data. We conducted our statistical analysis assuming that the values had been allocated normally. The chi-square tests (*Jin et al, 2006 [47]* and *Bahassine et al, 2018* [48)] are statistical tests. An explanation of the chi-square statistic along with the conditions under which the chi-square test is applied is provided below (*Jovic et al*, 2015) [35]:

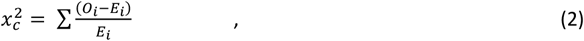

where c represents the degree of freedom, O represents the observed value, and E represents the expected value. By using a hand, you can find a fundamental chi-square value. For every single data item in your data set, you will have to perform a summation symbol calculation.

#### 3.3.2 Fisher score Feature Selection

According to Fisher’s score, a subset of the features should be selected that has a maximal distance between points from different classes should be large, while the distance between points within the same class should be minimized. The score of the i^th^ feature *S*_*i*_ will be calculated by Fisher Score *Q (Gu et al, 2012)* [50].

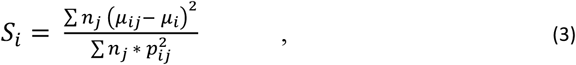

where *μ*_*ij*_ and *p*_*ij*_are the mean and variance of the i^th^ feature in the j^th^ class are considered, while *n*_*j*_ represents the number of instances in the j^th^ class, and *μ*_*i*_ denotes the mean of the i^th^ feature. In addition, it cannot handle redundant features. We argue that by implementing a generalized Fisher score, we can solve all these problems.

#### 3.3.3 Symmetric Uncertainty (SU) Feature Selection

The Symmetric Uncertain ty results in a more accurate way to select features defined entirely by class labels. Various research has used SU to evaluate the importance of features in classifying data (*Lin et al, 2019)* [51]. The definition of Symmetric Uncertainty (SU) is given by *Lin et al, 2019 [51]* and *Sosa-Cabrera et al, 2019* [52]:

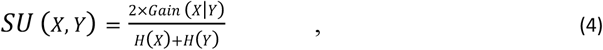

Here *H*(*X*)represents the entropy of variable *X*, and *Gain* (*X*|*Y*) denotes the information gain between variables *X* and *Y*, respectively. Assuming *P*(*X*) represents the prior probabilities for all possible values of *X*, the entropy of the discrete variable *X* is computed using Shannon’s formula as follows:

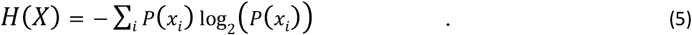

Entropy can also be defined conditionally on another variable, represented as *H*(*X*|*Y*). Given *P*(*X*), the prior probabilities of all possible values of *X*, and *P*(*X*|*Y*), (the posterior probabilities of *X* conditioned on *Y*), the conditional entropy of *X* given *Y* is expressed as:

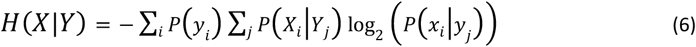

*Gain*(*X* |*Y*)quantifies the reduction in the uncertainty of *X* after observing *Y*, and is defined as follows:

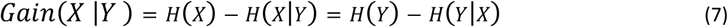

Information gain is a symmetrical measure and therefore *IG*(*X* |*Y*) = *IG*(*Y* |*X*). Information gain of two variables is normalized to the range 0,1 (that is SU). If two variables, *X* and *Y*, are independent, a totally dependent variable will have SU of 0, and a completely independent variable will have SU of 1. This means that if one variable’s value is predictable by the other, both variables will have SU of 1. Given *SU*(*X, Y*), which represents the symmetric uncertainty between variables *X* and *Y*, the T-Relevance refers to the relevance of a feature with respect to the target concept C, while the F-Correlation describes the relationship between two features. Specifically, the F-Correlation between any two distinct features*F*_*i*_ and *F*_*j*_ (*F*_*i*_ ∈ *F*_*j*_) ∧ (*i* ≠ *j*) is expressed as *SU* (*F*_*i*_, *F*_*j*_).

The relevance between a feature *F*_*i*_ (where *F*_*i*_ ∈ *F*_*j*_) and the target concept **C** is known as the T-Relevance of *F*_*i*_ with respect to **C**, and it is represented by *SU* (*F*_*i*_, *C*).

#### 3.3.4 Minimum Redundancy-Maximum Relevance (mRMR) Feature Selection

By using mRMR (Minimum Redundancy Maximal Relevance), features are selected that demonstrate a strong relationship with the classification while having low correlations themselves. In many feature selection algorithms, features and classification labels are evaluated with little attention to their mutual interactions. The minimum redundancy criterion is mentioned below (*Radovic et al, 2017 [53]* and *Seth et al, 2023)* [54]:

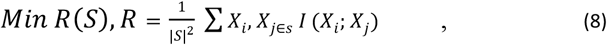

where *S* is the feature subset, |*S*| is the number of feature and ∑ *X*_*i*_; *X*_*j*_ is mutual information between feature i and j.

It measures relevance to pick out the value of mutual information. The circumstance for maximum relevance is described as follows.

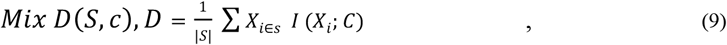

where *i* represents the mutual dependence involving the feature and *c*, which denotes the target activity. Based on the above redundancy and relevance measures, the mRMR feature selection algorithm is used. In accordance with condition (8) and (9), we fully considered both degrees of redundancy and degree of relevance when choosing features. This feature selection algorithm optimizes D and R at the same time since the following reason:

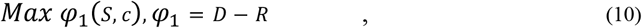

Or,

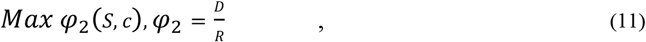

Where the feature subset *S*_*m*_ which is composed of m feature, according to (11) and (12) condition is the optimal (*m* + 1) feature from the feature subset *S* − *S*_*m*_, these two conditions are expressed as follows:

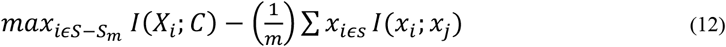

### 3.4 Used Classifiers

#### 3.4.1 Support Vector Machines (SVM)

An SVM analyses data samples accumulated over a period of time to locate a suitable hyperplane to segment them (*Taherzadeh et al, 2016)* [55]. In this study, a Linearly Separable Support Vector Machine is employed, where determining the margin within the sample space is essential, as the objective is to maximize it. The hyperplane is defined based on the following linear equation *(Taherzadeh et al, 2016 [55]* and *Tanveer et al, 2016* [56]):

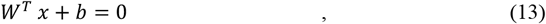

Here W is the normal vector that defines the orientation of the hyperplane, while b represents the offset, determining the measure of distance from the origin to the hyperplane. Consider the scenario where the hyperplane successfully classifies the training samples, i.e., the following formula applies for the training samples:

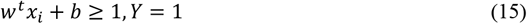

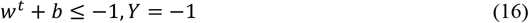

The above expression is referred to as the maximum margin hypothesis.

#### 3.4.2 Multi-Layer Perceptron

A MLPs is often called a “vanilla” neural network, particularly when it consists of just a single hidden layer. An MLP generally consists of a minimum of three layers of nodes: an input layer, a hidden layer, and an output layer. Each node, except those in the input layer, functions as a neuron that applies a nonlinear activation function (*Piccialli et al, 2022)* [57]. The error at output node *j* for the *n*^th^ data point can be expressed as followed by *e*_*j*_(*n*) = *d*_*j*_(*n*) − *y*_*j*_(*n*) In a perceptron, y represents the value produced by the perceptron and d is the target value. Using node weights as corrections, the overall output can be adjusted to minimize error, given by *Rosenblatt et al, 1961 [58]* and *Popescu et al, 2009* [59]:

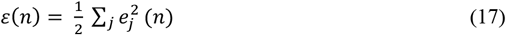

Using the gradient descent method, each weight is modified according to:

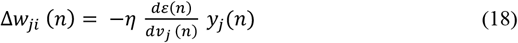

The learning rate, denoted by η, is selected to help the weights rapidly converge to a stable response without oscillations, where y refers to the output of the preceding neuron.

Induced local fields, /*nu*_*j*_, which themselves also vary, determine the derivative that must be calculated. A simple proof can be made that the derivative of an output node is described as follows:

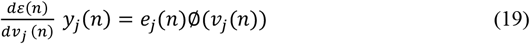

The derivative of ϕ corresponds to the activation function mentioned earlier, which remains constant. Modifying the weights connected to a hidden node involves deeper analysis, and the necessary derivative can be derived as follows:

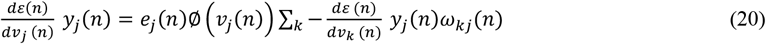

The output layer is influenced by the weights associated with the *k*-th node, reflecting the adjustment in weights.

### 3.5 Evaluation Metrics

All the aforesaid four feature selection methods were assessed in terms of three evaluation metrics, viz., accuracy, F1-score, and RMSE. The F1-score combines precision and recall into one metric by computing their harmonic mean. It is especially effective for evaluating and comparing the performance of two classifiers. RMSE represents the standard deviation of prediction errors that occur when a model is applied to a dataset. It resembles the Mean Squared Error (MSE); however, the square root of the MSE is calculated to evaluate the model’s accuracy.

## 4. Experimental Result and Discussion

In this part, the proposed technique is introduced with reference to its acquired consequences. Three case studies, namely, Microarray datasets from Mixed-Lineage Leukaemia’s (MLL), the central nervous system (CNS), lung cancer, ovarian cancer, and colon cancer are utilized to evaluate the accuracy of feature selection. The FS approach helps in filtering out irrelevant features from datasets with a high volume of insignificant data. By selecting a subset of features, FS aims to identify those that carry the most discriminative information. This process assists in choosing features (genes) with high relevance scores, while those with low relevance are discarded. Multiple datasets have been utilized to extract genes exhibiting high relevance scores.

### 4.1 Chi-Square feature selection with SVM and MLP classifier

The outcomes of the chi-square method have been verified using ‘microarray’ data across five datasets. The table displays statistical metrics, including the number of selected features, along with the corresponding accuracy, F1-score, and RMSE for each feature selection technique. It also highlights the accuracy achieved during both the training and testing stages. A detailed breakdown is provided in Table 2 and Table 3, summarizing the statistics for five datasets: Mixed-Lineage Leukemia (MLL), Central Nervous System, Lung Cancer, Ovarian Cancer, and Colon Cancer, respectively.

**Table 2.** Performance analysis of the SVM classifier employing chi-square for feature selection across five datasets.

| **Accuracy (%)** | **F1-score** | **RMSE** | **Dataset Name** |
| --- | --- | --- | --- |
| 91.42 | 0.909 | 0.213 | Mixed lineage leukemias (MLL) |
| 84.82 | 0.776 | 0.324 | colon cancer |
| 89.30 | 0.889 | 0.311 | central nervous system (CNS) |
| 94.40 | 0.910 | 0.201 | Lung Cancer |
| 93.90 | 0.926 | 0.218 | Ovarian Cancer |

**Table 3.** Performance analysis of the MLP classifier employing chi-square for feature selection across five datasets.

| **Accuracy (%)** | **F1-score** | **RMSE** | **Dataset Name** |
| --- | --- | --- | --- |
| 86.40 | 0.828 | 0.172 | Mixed-lineage leukaemia's (MLL) |
| 82.31 | 0.766 | 0.393 | colon cancer |
| 66.67 | 0.657 | 0.411 | central nervous system (CNS) |
| 97.56 | 0.969 | 0.145 | Lung Cancer |
| 88.24 | 0.878 | 0.198 | Ovarian Cancer |

From **Table 2**, the SVM classifier with chi-square feature selection achieved 91.42% accuracy, a 0.909 F1-score, and 0.213 RMSE on the MLL dataset using six features. For the Colon Cancer dataset, it obtained 84.82% accuracy, a 0.776 F1-score, and 0.324 RMSE. On the CNS dataset, the results were 89.30% accuracy, a 0.889 F1-score, and 0.311 RMSE. The best performance was recorded for Lung Cancer and Ovarian Cancer datasets with accuracies of 94.40% and 93.90%, highlighting the effectiveness of the chi-square method. From **Table 3**, the MLP classifier with an alternative feature selection method achieved 86.40% accuracy, a 0.828 F1-score, and 0.172 RMSE on the MLL dataset. For Colon Cancer, it obtained 82.31% accuracy, a 0.766 F1-score, and 0.393 RMSE. Performance on the CNS dataset was lower, with 66.67% accuracy, a 0.657 F1-score, and 0.411 RMSE. The highest accuracies were recorded for Lung Cancer (97.56%) and Ovarian Cancer (88.24%), demonstrating the method’s effectiveness.

### 4.2 Fisher score Feature selection with SVM and MLP classifier

The performance of the Fisher score method has been validated using ‘microarray’ data across five datasets. The table presents various metrics, including the number of selected features, along with the achieved accuracy, F1-score, and RMSE for each feature selection method, as well as the accuracy obtained during both the training and testing phases. A detailed analysis is provided in Table 4 and Table 5, which present the statistics for five datasets: Mixed-Lineage Leukaemia (MLL), Central Nervous System, Lung Cancer, Ovarian Cancer, and Colon Cancer, respectively.

**Table 4.** Performance of the SVM classifier with Fisher score as a feature selection method with five datasets.

| Accuracy (%) | F1-score | RMSE | Dataset Name |
| --- | --- | --- | --- |
| 94.28 | 0.938 | 0.129 | Mixed-lineage leukaemia's (MLL) |
| 84.28 | 0.797 | 0.348 | colon cancer |
| 83.33 | 0.833 | 0.201 | central nervous system (CNS) |
| 95.12 | 0.951 | 0.049 | Lung Cancer |
| 96.42 | 0.964 | 0.036 | Ovarian Cancer |

**Table 5.** Performance of the MLP classifier with Fisher score as a feature selection method with five datasets.

| Accuracy (%) | F1-score | RMSE | Dataset Name |
| --- | --- | --- | --- |
| 81.34 | 0.805 | 0.235 | Mixed-lineage leukaemia's (MLL) |
| 76.59 | 0.765 | 0.377 | colon cancer |
| 75.00 | 0.750 | 0.311 | central nervous system (CNS) |
| 92.68 | 0.926 | 0.180 | Lung Cancer |
| 90.56 | 0.905 | 0.195 | Ovarian Cancer |

**Table 4** shows that the SVM classifier with optimized feature selection achieved 94.28% accuracy, a 0.938 F1-score, and 0.129 RMSE on the MLL dataset. For Colon Cancer, it reached 84.28% accuracy, a 0.797 F1-score, and 0.348 RMSE. The CNS dataset achieved 83.33% accuracy, a 0.833 F1-score, and 0.201 RMSE. Lung and Ovarian Cancer datasets showed the best results with 95.12% and 96.42% accuracies and very low RMSEs (0.049 and 0.036), confirming the effectiveness of the optimized feature selection method. **Table 5** shows that the SVM classifier with a different feature selection method achieved 81.34% accuracy, a 0.805 F1-score, and 0.235 RMSE on the MLL dataset. For Colon Cancer, it obtained 76.59% accuracy, a 0.765 F1-score, and 0.377 RMSE, while the CNS dataset recorded 75.00% accuracy, a 0.750 F1-score, and 0.311 RMSE. Meanwhile, the Lung Cancer and Ovarian Cancer datasets exhibited the highest performance, with accuracies of 92.68% and 90.56%, respectively, and lower RMSE values of 0.180 and 0.195. These results indicate that while the feature selection method is effective, its performance varies across different datasets.

### 4.3 Symmetrical Uncertainty (SU) Feature selection with SVM and MLP classifier

The SU results were validated using microarray data across five datasets. **Tables 6** and **7** present the number of selected features along with accuracy, F1-score, and RMSE for each feature selection method during training and testing. The datasets include MLL, CNS, Lung Cancer, Ovarian Cancer, and Colon Cancer.

**Table 6.** Performance of the SVM classifier with Symmetrical Uncertainty (SU) as a feature selection method with five datasets.

| Accuracy (%) | F1-score | RMSE | Dataset Name |
| --- | --- | --- | --- |
| 67.85 | 0.618 | 0.616 | Mixed-lineage leukaemia's (MLL) |
| 74.52 | 0.630 | 0.460 | colon cancer |
| 76.00 | 0.760 | 0.340 | central nervous system (CNS) |
| 95.23 | 0.952 | 0.148 | Lung Cancer |
| 96.08 | 0.960 | 0.140 | Ovarian Cancer |

From **Table 6**, It is observed that the SVM classifier with an alternative feature selection method achieved 67.85% accuracy, a 0.618 F1-score, and 0.616 RMSE on the MLL dataset. The Colon Cancer dataset performed slightly better with 74.52% accuracy. The CNS dataset reached 76.00% accuracy, while Lung and Ovarian Cancer datasets showed the highest performance with 95.23% and 96.08% accuracy, and low RMSE values of 0.148 and 0.140, respectively. This highlight varying effectiveness across datasets. From **Table 7**, The SVM classifier with a different feature selection method achieved 70.80% accuracy on MLL, 71.75% on Colon Cancer, and 66.67% on CNS datasets. Lung Cancer showed the best performance with 92.68% accuracy, while Ovarian Cancer reached 74.51%. The results highlight strong performance for Lung Cancer but relatively lower effectiveness for MLL and CNS datasets.

**Table 7.** Performance of the MLP classifier with Symmetrical Uncertainty (SU) as a feature selection method with five datasets.

| Accuracy (%) | F1-score | RMSE | Dataset Name |
| --- | --- | --- | --- |
| 70.80 | 0.789 | 0.704 | Mixed-lineage leukaemia's (MLL) |
| 71.75 | 0.717 | 0.313 | colon cancer |
| 66.67 | 0.666 | 0.334 | central nervous system (CNS) |
| 92.68 | 0.926 | 0.174 | Lung Cancer |
| 74.51 | 0.745 | 0.255 | Ovarian Cancer |

### 4.4 Minimum redundancy-maximum relevance (mRMR) Feature selection with SVM and MLP classifier

The results of mRMR were validated using microarray data for two datasets. The tables present statistics such as the number of features selected, accuracy, F1-score, and RMSE for each FS method during training and testing. Specifically, **Tables 8-9** summarize the results for five datasets: MLL, CNS, Lung Cancer, Ovarian Cancer, and Colon Cancer.

**Table 8.** Performance of the SVM classifier with (mRMR) as a feature selection method with five datasets.

| Accuracy (%) | F1-score | RMSE | Dataset Name |
| --- | --- | --- | --- |
| 94.28 | 0.937 | 0.188 | Mixed-lineage leukaemia's (MLL) |
| 88.80 | 0.867 | 0.279 | colon cancer |
| 67.67 | 0.667 | 0.333 | central nervous system (CNS) |
| 75.61 | 0.756 | 0.244 | Lung Cancer |
| 100.00 | 1.000 | 0.000 | Ovarian Cancer |

**Table 9.** Performance of the MLP classifier with (mRMR) as a feature selection method with five datasets.

| Accuracy (%) | F1-score | RMSE | Dataset Name |
| --- | --- | --- | --- |
| 88.84 | 0.896 | 0.170 | Mixed-lineage leukaemia's (MLL) |
| 79.56 | 0.796 | 0.218 | colon cancer |
| 58.33 | 0.583 | 0.417 | central nervous system (CNS) |
| 85.37 | 0.853 | 0.247 | Lung Cancer |
| 100.00 | 1.000 | 0.001 | Ovarian Cancer |

**Table 8** shows that the SVM classifier with a specific feature selection method achieved 94.28% accuracy, a 0.937 F1-score, and an RMSE of 0.188 for the MLL dataset. For Colon Cancer, the accuracy was 88.80%, while CNS showed lower performance with 67.67% accuracy. Lung Cancer achieved 75.61% accuracy. Remarkably, Ovarian Cancer achieved perfect results with 100% accuracy and an RMSE of 0.000, highlighting the method’s strong effectiveness on certain datasets. From **Table 9**, using an advanced feature selection method, the SVM classifier achieved 88.84% accuracy, a 0.896 F1-score, and a RMSE of 0.170 on the MLL dataset. For Colon Cancer, the accuracy was 79.56%, while CNS showed the lowest performance at 58.33%. Lung Cancer achieved 85.37% accuracy. Notably, Ovarian Cancer reached 100% accuracy with an almost zero RMSE, highlighting the method’s strong effectiveness on simpler datasets but the need for improvement on more complex ones like CNS.

### 4.5 Support vector machine

The Support Vector Machine (SVM) classifier was evaluated using various feature selection methods across five datasets, with the resulting accuracies summarized in **Table 10**. In **Table 10**, it is evident when mRMR is used as a feature selection method, it achieves a perfect accuracy of 100.00%, an F1-score of 1.000, and an RMSE of 0.000 for the Ovarian Cancer dataset. Similarly, Fisher Score and Symmetrical Uncertainty (SU) also demonstrate high performance on the Ovarian Cancer dataset, with accuracy values of 96.42% and 96.08%, respectively. The Chi-square method attained 94.40% accuracy, a 0.910 F1-score, and a 0.201 RMSE for the Lung Cancer dataset, highlighting the strong impact of feature selection on model performance across datasets.

**Table 10.** SVM Performance Using Different Feature Selection Techniques.

| Feature selection method | Accuracy (%) | F1-score | RMSE | Dataset |
| --- | --- | --- | --- | --- |
| Chi-square | 94.40 | 0.910 | 0.201 | Lung Cancer |
| Fisher Score | 96.42 | 0.964 | 0.036 | Ovarian Cancer |
| Symmetrical Uncertainty (SU) | 96.08 | 0.960 | 0.140 | Ovarian Cancer |
| mRMR | 100.00 | 1.000 | 0.000 | Ovarian Cancer |

### 4.6 Multi-layer Perceptron

The Multi-layer Perceptron (MLP) classifier was evaluated using various feature selection methods on two datasets, with the resulting accuracies summarized in **Table 11**. As shown in **Table 11**, using mRMR for feature selection leads to perfect classification on the Ovarian Cancer dataset, achieving 100% accuracy, an F1-score of 1.000, and an RMSE of 0.001. For the Lung Cancer dataset, the Chi-square method delivers the best results with 97.56% accuracy, an F1-score of 0.969, and an RMSE of 0.145. Additionally, Fisher Score and Symmetrical Uncertainty (SU) both achieve 92.68% accuracy, with similar F1-scores and RMSE values. These findings demonstrate how different feature selection methods affect model performance across datasets.

**Table 11.**
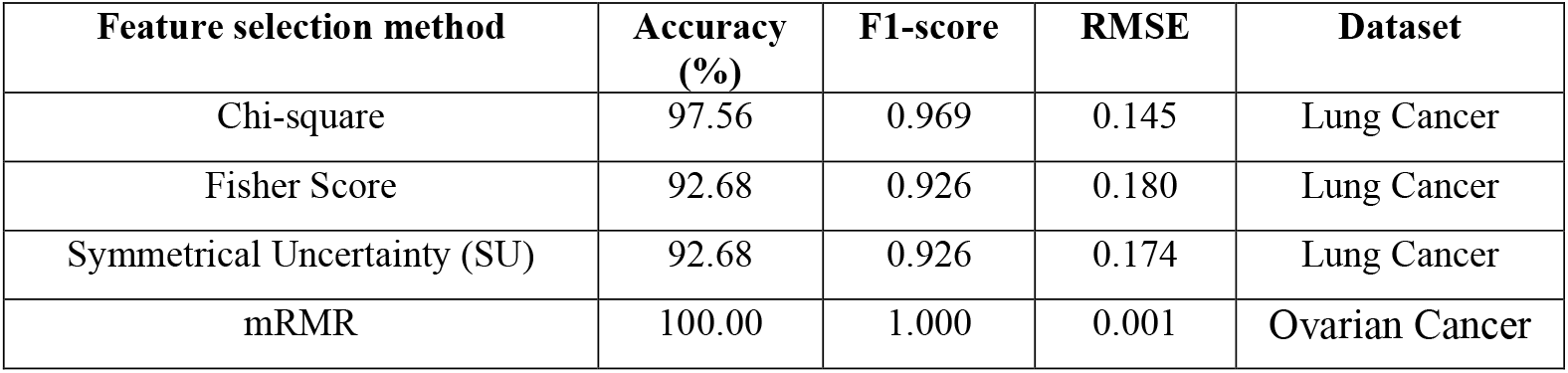
Performance of MLP Using Different Feature Selection Techniques.

## 5. CONCLUSION

In this research article, the main objective of this work was to thoroughly investigate well-known filter-based feature selection techniques utilizing univariate and multivariate analysis. Four leading filter techniques were carefully analysed across five microarray datasets to deliver a comparative evaluation of their effectiveness. Our analysis revealed that among the four feature selection methods, Maximum Relevance Minimum Redundancy (mRMR) consistently demonstrated superior performance. It outperformed the others in terms of average accuracy, F1-score, and Root Mean Square Error (RMSE), signifying its effectiveness in identifying the most informative features. Additionally, the Fisher Score, Chi-square, and Symmetrical Uncertainty (SU) methods exhibited commendable results, securing their positions as the second, third, and fourth best methods, respectively. While they did not surpass the mRMR method, their contributions to enhancing classification outcomes are noteworthy and warrant consideration in various gene expression data analysis scenarios.

In summary, this study offers important insights into the comparative strengths of various feature selection techniques when applied to microarray datasets. The outstanding performance of the mRMR method, along with the solid results from other methods, highlights the critical role of careful feature selection in maximizing the utility of high-dimensional biological data. Our findings contribute to the advancement of bioinformatics and lay the groundwork for future research focused on improving gene expression analysis for key applications like disease diagnosis and tumor classification.

**Figure 1:**
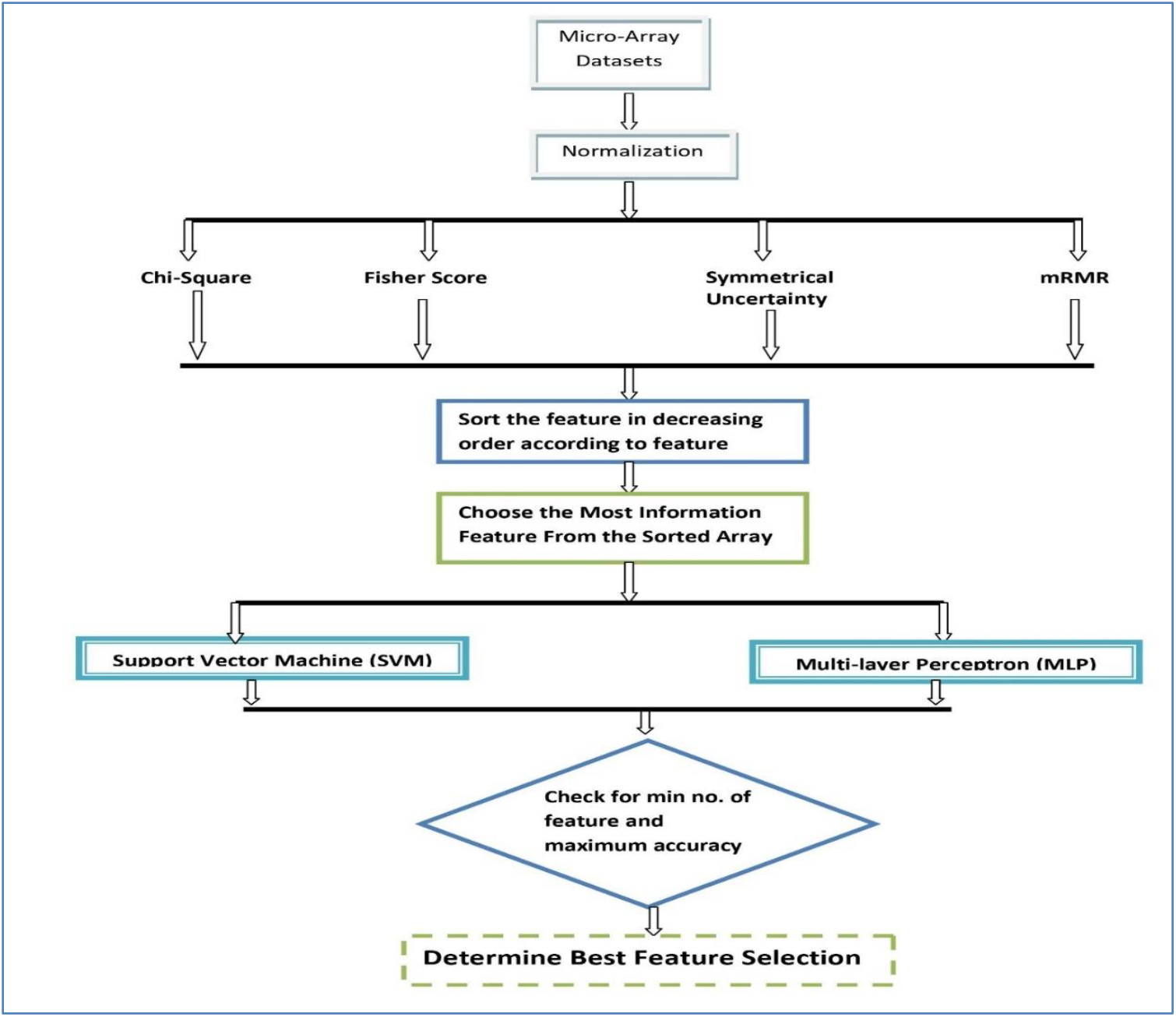
Experimental Framework.

**Figure 2:**
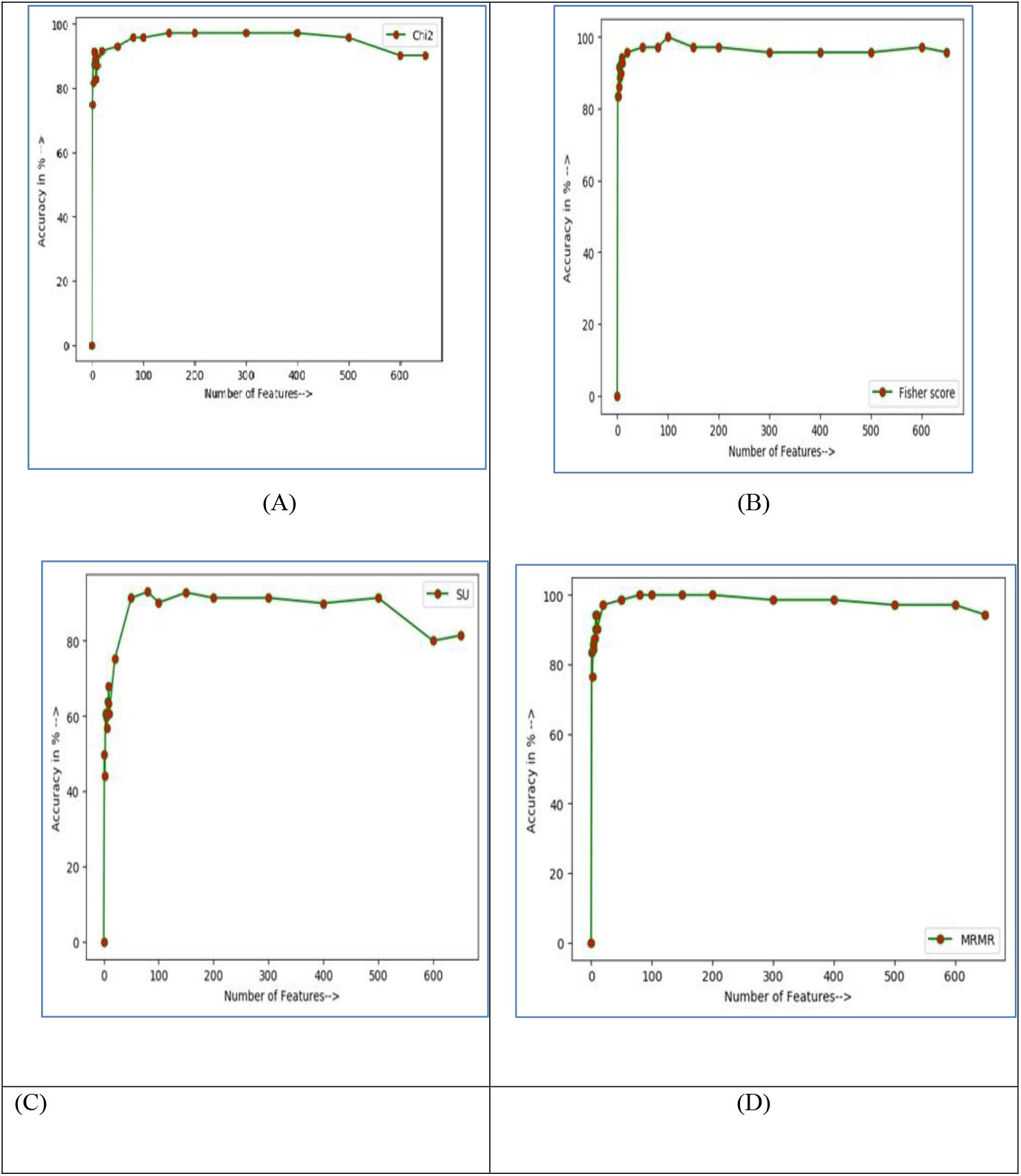
Increasing number of features vs. accuracy graph for (A) chi-square, (B) Fisher Score, (C) Symmetrical Uncertainty (SU), (D) Minimum redundancy maximum relevance (mRMR) feature selection method.

## Declarations

### Ethics approval and consent to participate

Not applicable.

### Consent for publication

Not applicable.

### Institutional Review Board Statement

Not applicable.

### Research involving Human Participants and/or Animals

This work doesn’t have any connection with human involvement or animal involvement.

### Informed Consent Statement

Not applicable.

### Conflicts of Interest

The authors declare no conflict of interest.

### Authors’ Contributions

A.I., T.B., K.M. performed the experiments. A.I., T.B., K.M., K.A., S.M. have done the validation and assessment of the models and written the manuscript. R.K.B., J.G., S.M. supervised the work and helped with manuscript editing.

### Funding

The authors received no funding from their institutes.

### Data Availability

The datasets generated and/or analyzed during the current study are available in the Mixed-lineage leukaemia’s (MLL), central nervous system, Lung Cancer, Ovarian Cancer and colon cancer datasets, https://csse.szu.edu.cn/staff/zhuzx/datasets.html [34].

### Author Biographies

**Atikul Islam** is currently working as an asst. professor in the department of Computer Science and Engineering, Seacom Engineering College, Sankrail, Howrah, India. He is also pursuing in the department of Computer Science and Engineering, Kalyani University, Kalyani, India. His research interests include bioinformatics and machine learning.

**Tapas Bhadra** is currently working as an asst. professor in Department of Computer Science and Engineering, Aliah University, Kolkata, West Bengal, India. His research interests include feature mining and machine learning.

**Kalyani Mali** is currently working as an asst. professor in the department of Computer Science and Engineering, University of Kalyani, Kalyani, India. His research interests include Pattern Recognition, Image Processing, Data Mining, Soft Computing.

**Jayant Giri** is an asst prof. in the Department of Mechanical Engineering, Yeshwantrao Chavan College of Engineering, Nagpur, India. His research interests include optimization, modelling and simulation.

**Khursheed Aurangzeb** is an assistant professor in the department of Computer Engineering, College of Computer and Information Sciences, King Saud University, Riyadh, Saudi Arabia. His research interests are data mining and deep Learning.

**Ram Kaji Budhathoki** is an associate professor in the department of Electronics Engineering, Kathmandu University, Dhulikhel, Nepal. His research interests are electrical engineering, biosensor, feature mining.

**Saurav Mallik** is currently working as a research scientist, dept of Pharmacology & Toxicology, University of Arizona, Tucson, USA. He previously worked as a postdoctoral fellow at Harvard University, Boston, USA. His research interests are computational Biology, data mining.

## References

[1] Bolón-Canedo, V., Sánchez-Maroño, N., Alonso-Betanzos, A., Benítez, J. M., Herrera, F., “A review of micro-array data sets and applied feature selection methods”, information sciences, Volume 282, Issue 6, 111–135, 2014.

[2] C. Lazar et al, “A Survey on Filter Techniques for Feature Selection in Gene Expression Microarray Analysis”, in IEEE/ACM Transactions on Computational Biology and Bioinformatics, Volume 9, Issue 4, 1106–1119, 2012.

[3] Kohavi R. and John G., “Wrappers for Feature Subset Selection”, Artificial Intelligence, Volume 97, 273–324, 1997.

[4] Hanai, T., Hamada, H., Okamoto, M., “Application of bioinformatics for DNA microarray data to bioscience,bioengineering and medical fields”, Journal of Bioscience and Bioengineering, Volume 101, Issue 5, 377–384, 2006.

[5] Park, T., Yi, SG., Kang, SH. et al., “Evaluation of normalization methods for microarray data”, BMC Bioinformatics, Volume 4, Issue 33, 2003.

[6] Liu, X., Li, N., Liu, S., Wang, J., Zhang, N., Zheng, X., et al., “Normalization Methods for the Analysis of Unbalanced Transcriptome Data: A Review.”, Frontiers in Bioengineering and Biotechnology, Volume 7, 2019.

[7] Singh, D., Singh, B., “Investigating the impact of data normalization on classification performance”, Applied Soft Computing, 2019.

[8] S. Gopal Krishna Patro, Kishore Kumar sahu, “Normalization: A Preprocessing Stage”, ieee, Volume 102, 2015.

[9] Hira, Z. M., Gillies, D. F., “A Review of Feature Selection and Feature Extraction Methods Applied on Microarray Data”, Advances in Bioinformatics, Volume 1, 1–13, 2015.

[10] Emura, T., Matsui, S., Chen, H.-Y. (2018). compound.Cox: univariate feature selection and compound covariate for predicting survival. Computer Methods and Programs in Biomedicine, 2018.

[11] Sumaiya Thaseen, I., Aswani Kumar, C. “Intrusion detection model using fusion of chi-square feature selection and multi class SVM”, Journal of King Saud University Computer and Information Sciences, 29(4), 462–472, 2019

[12] Bahassine, S., Madani, A., Al-Sarem, M., Kissi, M., “Feature selection using an improved Chi-square for Arabic text classification”, Journal of King Saud University - Computer and Information Sciences., 2018.

[13] Sun, L., Wang, T., Ding, W., Xu, J., Lin, Y. “Feature selection using Fisher score and multilabel neighborhood rough sets for multilabel classification”. Information Sciences, 578, 887–912, 2021.

[14] Quanquan Gu and Zhenhui Li and Jiawei Han. “Generalized Fisher Score for Feature Selection”, IEEE, 2012.

[15] Sun, Y., Li, J., Liu, J. et al. “Using causal discovery for feature selection in multivariate numerical time series”. Mach Learn, 101, 377–395, 2015.

[16] Senthamarai Kannan, S., Ramaraj, N. “A novel hybrid feature selection via Symmetrical Uncertainty ranking based local memetic search algorithm”, Knowledge-Based Systems, 23(6), 580–585, 2010.

[17] Lin, X., Li, C., Ren, W., Luo, X., Qi, Y. “A new feature selection method based on symmetrical uncertainty and interaction gain”, Computational Biology and Chemistry, 107149, 2019.

[18] Sai Prasad Potharaju and M. Sreedevi. “A Novel M-Cluster of Feature Selection Approach Based on Symmetrical Uncertainty for Increasing Classification Accuracy of Medical Datasets”, JOURNAL OF Engineering Science and Technology Review, 2017.

[19] De Jay, N., Papillon-Cavanagh, S., Olsen, C., El-Hachem, N., Bontempi, G., Haibe-Kains, B. “mRMRe: an R package for parallelized mRMR ensemble feature selection”. Bioinformatics, 29(18), 2365–2368, 2013.

[20] Li, B.-Q., Hu, L.-L., Chen, L., Feng, K.-Y., Cai, Y.-D., Chou, K.-C. “Prediction of Protein Domain with mRMR Feature Selection and Analysis”. PLoS ONE, 7(6), e39308, 2012.

[21] Furey, T. S., Cristianini, N., Duffy, N., Bednarski, D. W., Schummer, M., Haussler, D. “Support vector machine classification and validation of cancer tissue samples using microarray expression data”, Bioinformatics, 16(10), 906–914, 2000.

[22] Gold, C., Sollich, P., “Model selection for support vector machine classification”. Neurocomputing, 55(1-2), 221–249, 2003.

[23] Zhang, Y., “Support Vector Machine Classification Algorithm and Its Application”. Information Computing and Applications, 179–186, 2012.

[24] Zhang, J., Liu, S., Wang, Y. “Gene association study with SVM, MLP and cross-validation for the diagnosis of diseases”. Progress in Natural Science, 18(6), 741–750, 2008.

[25] Khalsan, Mahmood & Mu, M. & Al-Shamery, Eman & Machado, Lee & Ajit, Suraj & Opoku Agyeman, Michael. (2023). Fuzzy Gene Selection and Cancer Classification Based on Deep Learning Model. 10.48550/arXiv.2305.04883.

[26] Khan, Zardad & Ali, Amjad & Aldahmani, Saeed. (2024). Feature Selection via Robust Weighted Score for High Dimensional Binary Class-Imbalanced Gene Expression Data. Heliyon. 10. e38547. 10.1016/j.heliyon.2024.e38547.

[27] Nematzadeh, Hossein & Mani, Joseph & Nematzadeh, Zahra & Akbari, Ebrahim & Mohamad, Radziah. (2025). Distance-based mutual congestion feature selection with genetic algorithm for high-dimensional medical datasets. Neural Computing and Applications. 37. 6217–6232. 10.1007/s00521-024-10837-4.

[28] A. Osareh and B. Shadgar, “Machine learning techniques to diagnose breast cancer,”, 2010 5th International Symposium on Health Informatics and Bioinformatics, 2010.

[29] Salem DA, Seoud A, Ahmed R, Ali HA. “MGS-CM: a multiple scoring gene selection technique for cancer classification using microarrays”. Int J Comput Appl, 2011.

[30] Wang L, Chu F, Xie W. Accurate cancer classification using expressions of very few genes. IEEE/ACM Trans Comput Biol Bioinformatics (TCBB) 2007.

[31] Zhang J-G, Deng H-W. Gene selection for classification of microarray data based on the Bayes error. BMC Bioinformat 2007;8(1):370–8.

[32] X. Hang, “Cancer classification by sparse representation using microarray gene expression data”, 2008 IEEE International Conference on Bioinformatics and Biomeidcine Workshops, 2008, pp. 174–177.

[33] Bharathi A, Natarajan A. “Cancer classification of bioinformatics data using ANOVA”. Int J Comput Theory Eng 2010;2(3):369–73.

[34] Tang K-l, Yao W-j, Li T-h, Li Y-x, Cao Z-W. “Cancer classification from the gene expression profiles by discriminant Kernel-PLS”. J Bioinformat Comput Biol 2010;8(Suppl. 01):147–60.

[35] Furey TS, Cristianini N, Duffy N, Bednarski DW, Schummer M, Haussler D. “Support vector machine classification and validation of cancer tissue samples using microarray expression data”. Bioinformatics 2000;16(10):906–14.

[36] Guyon I, Weston J, Barnhill S, Vapnik V. “Gene selection for cancer classification using support vector machines”. Mach Learn 2002;46(1–3):389–422.

[37] Ye J, Li T, Xiong T, Janardan R. “Using uncorrelated discriminant analysis for tissue classification with gene expression data”. IEEE/ACM Trans Comput Biol Bioinformat (TCBB) 2004;1(4):181–90.

[38] Liu JJ, Cutler G, Li W, Pan Z, Peng S, Hoey T, et al. “Multiclass cancer classification and biomarker discovery using GA-based algorithms”. Bioinformatics 2005;21(11):2691–7.

[39] Yu L, Liu H. “Redundancy based feature selection for microarray data”. In: Proceedings of the tenth ACM SIGKDD international conference on knowledge discovery and data mining. New York: ACM; 2004. p. 737–42.

[40] Peng Y, Li W, Liu Y. “A hybrid approach for biomarker discovery from microarray gene expression data for cancer classification”. Cancer Informat 2007;2:301–11.

[41] Liu, X., Li, N., Liu, S., Wang, J., Zhang, N., Zheng, X., et al. “Normalization Methods for the Analysis of Unbalanced Transcriptome Data: A Review.”, Frontiers in Bioengineering and Biotechnology, Volume 7, 2019.

[42] Singh, D., Singh, B., “Investigating the impact of data normalization on classification performance”, Applied Soft Computing, 2019.

[43] Islam, A., Seth, S., Bhadra, T., Mallik, S., Roy, A., Li, A., & Sarkar, M. (2024). Feature Selection, Clustering, and IoMT on Biomedical Engineering for COVID-19 Pandemic: A Comprehensive Review. Journal of Data Science and Intelligent Systems, 2(4), 191–204. 10.47852/bonviewJDSIS3202916

[44] Li J, Liu H. Kent ridge bio-medical dataset. http://datam.i2r.a-star.edu.sg/datasets/krbd/index.html [access date: 09 August 2024].

[45] Zexuan Zhu, Y. S. Ong and M. Dash, “Markov Blanket-Embedded Genetic Algorithm for Gene Selection”, Pattern Recognition, Vol. 49, No. 11, 3236–3248, 2007

[46] Golub TR, Slonim DK, Tamayo P, Huard C, Gaasenbeek M, Mesirov JP, et al. Molecular classification of cancer: class discovery and class prediction by gene expression monitoring. Science 1999;286(5439):531–7.

[47] Jin, X., Xu, A., Bie, R., Guo, P., “Machine Learning Techniques and Chi-Square Feature Selection for Cancer Classification Using SAGE Gene Expression Profiles.”, Data Mining for Biomedical Applications, 106–115, 2006.

[48] Bahassine, S., Madani, A., Al-Sarem, M., Kissi, M., “Feature selection using an improved Chi-square for Arabic text classification”, Journal of King Saud University - Computer and Information Sciences., 2018.

[49] Jovic, A., Brkic, K., Bogunovic, N., “A review of feature selection methods with applications”, International Convention on Information and Communication Technology, Electronics and Microelectronics (MIPRO), 2015.

[50] Q. Gu,Zhenhui Li,J. Han et al., “Generalized Fisher Score for Feature Selection”, in IEEE/ACM Transactions on Computational Biology and Bioinformatics, 2012.

[51] Lin, X., Li, C., Ren, W., Luo, X., Qi, Y., “A new feature selection method based on symmetrical uncertainty and interaction gain”, Computational Biology and Chemistry, 2019.

[52] Sosa-Cabrera, G., García-Torres, M., Gómez-Guerrero, S., Schaerer, C. E., Divina, F., “A Multivariate approach to the Symmetrical Uncertainty Measure: Application to Feature Selection Problem”, Information Sciences, 2019.

[53] Radovic, M., Ghalwash, M., Filipovic, N. et al. “Minimum redundancy maximum relevance feature selection approach for temporal gene expression data”, BMC Bioinformatics, Volume 18, Issue 9, 2017.

[54] Seth, S., Mallik, S., Islam, A., Bhadra, T., Roy, A., Singh, P. K., … & Zhao, Z. (2023). Identifying genetic signatures from single-cell RNA sequencing data by matrix imputation and reduced set gene clustering. Mathematics, 11(20), 4315.

[55] G. Taherzadeh, Y. Zhou, A. W.-C. Liew, and Y. Yang, “Sequence-based prediction of protein-carbohydrate binding sites using support vector machinest”, Journal of Chemical Information and Modeling, vol. 56, no. 10, pp. 2115–2122, 2016.

[56] M. Tanveer, M. A. Khan, and S.-S. Ho, “Robust energy-based least squares twin support vector machines”, Applied Intelligence, vol. 45, no. 1, pp. 174–186, 2016.

[57] Piccialli, V., Sciandrone, M. Nonlinear optimization and support vector machines. Ann Oper Res 314, 15–47 (2022). 10.1007/s10479-022-04655-x

[58] Rosenblatt, Frank. x. “Principles of Neurodynamics: Perceptrons and the Theory of Brain Mechanisms”. Spartan Books, Washington DC, 1961.

[59] Popescu, Marius-Constantin & Balas, Valentina & Perescu-Popescu, Liliana & Mastorakis, Nikos. (2009). Multilayer perceptron and neural networks. WSEAS Transactions on Circuits and Systems. 8.

